# Predation impacts late but not early community assembly in model marine biofilms

**DOI:** 10.1101/2020.08.06.233379

**Authors:** Sven P. Tobias-Hünefeldt, Jess Wenley, Federico Baltar, Sergio E. Morales

**Author notes:** Correspondance: Dr. Sergio E. Morales, University of Otago, Department of Microbiology and Immunology, 720 Cumberland Street, North Dunedin, Dunedin 9054, New Zealand.

## Abstract

Bottom-up selection plays an important role in microbial community assembly but is unable to account for all observed variance. Other processes like top-down selection (e.g. predation) may be partially responsible for the unexplained variance. However, top-down processes often remain unexplored, especially in interaction with bottom-up selective pressures. We utilised an *in situ* marine biofilm model system to test the effects of bottom-up (i.e. substrate properties) and top-down (i.e. predator exclusion via 100 µm mesh) selective pressures on community assembly over time (56 days). Community compositions were monitored using 16S and 18S rRNA amplicon sequencing. Wooden substrates promoted heterotrophic growth, while the inert substrates’ (i.e., plastic, glass, tile) lack of degradable material selected for autotrophs. Early wood communities contained 9-50% more mixotrophs and heterotrophs (e.g. Proteobacteria and Euglenozoa) compared to inert substrates. Inert substrates instead showed twice the autotrophic (e.g. Cyanobacteria and Ochrophyta) abundance. Late communities differed mainly due to exclusion status, as large predators preferably pruned heterotrophs. This resulted in the autotrophic domination of native communities, while high heterotrophic abundance characterised exclusive conditions. Top-down control through exclusion increased explainable variance by 18-53%, depending on community age, leading to increased understanding of the underlying ecological framework that guides microbial community assembly.

## Introduction

Sequencing technology advances have increased insight into microbiome compositions across all ecosystems, but mechanistic understanding of community succession remains poor. Historic pure culture approaches, which remove biotic interactions, have led to a focus on resource limitations as a mechanism to explain differences in growth and community composition^1,2^. However, biotic interactions have the potential to impact microbiome assembly and composition^3^. Microbiome studies generally report 20-67%^4–6^ unexplained composition variance when utilising solely resource limitations as a selective pressure. Unexplained variance represents a fundamental knowledge gap within the microbial ecology field and suggests that other processes must be contributing to microbiome changes.

Previous studies have found that communities can be shaped by two possible overarching processes: stochastic and deterministic. A stochastic process increases a community’s inherent randomness, producing variable microbiomes under identical conditions^7^. Early communities are commonly stochastic^8^, the inherent randomness of dispersal and ecological drift driving composition^9,10^. In contrast, deterministic processes, focused upon in the present study, shape communities by selecting for or against specific organisms^11–13^. These complementary processes shape structure by representing extremes along the same continuum, and a process such as dispersal may be equally stochastic and deterministic in a context and habitat-dependent manner^9^. Dispersal rates are stochastic if they rely on population size, but species traits and their activity level mean that dispersal can also be deterministic^9^.

Both abiotic and biotic deterministic processes can influence community assembly. Abiotic factors are chemical or physical environmental components (such as nutrients, pH, temperature, pressure) that affect community composition and function^14–16^. Meanwhile biotic factors are defined as selective pressures related to, or resulting from, living organisms that may affect community composition. Biotic factors can concern the biotic interactions between organisms, and these interactions can result in the promotion or inhibition of organism abundance. The cooperation between microbes^17,18^, and microbes and higher trophic levels (i.e. larger predators) increases both participants’ abundance. One such example is a Copepods ability to affect environmental community compositions by farming their own microbiome^19^. Meanwhile predation promotes the growth of one organism at the expense of another^20,21^. Both abiotic^22^ and biotic^23^ factors are dynamic throughout space and time. These shifting environmental conditions and biotic interactions change the community over time due to continuous selective pressures^22,24,25^.

Selective pressures enact compositional changes through autotrophs (i.e. primary producers) in a bottom-up dependent manner^26^, or through predators in a top-down dependent manner^27^. Previous studies on environmental microbiology have typically focused on bottom-up controls^4,28^. Field studies concerning top-down controls are rarer, and bottom-up and top-down selective pressure interaction effects on microbial communities remain largely unexplored^29–34^. Bottom-up microbiome studies commonly report unexplained variance, which could be partially driven by top-down controls^35,36^. Identifying the contribution of top-down controls to total variance is crucial to better understand community assembly mechanisms. Therefore, our study focuses on comparing the effects of bottom-up (i.e. different substrate surface) and top-down (i.e. predator presence and exclusion via a mesh enclosure) on compositional variance within marine biofilms.

We aim to quantify the influence of top-down and bottom-up controls on community assembly in a model marine biofilm system. We hypothesise that bottom-up controls drive early assembly, while established communities are controlled equally by both bottom-up and top-down mechanisms. To test these hypotheses, we created a model biofilm system using four different substrates (i.e. plastic, glass, tile, wood) to represent a degradability and surface property gradient and thus bottom-up selection. Biofilms were reared *in situ* under native (non-enclosed) or enclosed (exclusion with 100 µm mesh) conditions to model top-down influence by limiting predator access. Prokaryotic and eukaryotic community successions and composition were monitored via 16S and 18S rRNA amplicon sequencing over a 56-day period. We used here an *in situ* marine biofilm as a model system to study the integration of both bottom-up and top-down selective pressures on microbial communities assembly. Yet, the knowledge derived from this experiment would be applicable across all ecosystems, as we assess the underlying framework rather than habitat specific processes.

## Methods

### Sample preparation and collection

Individual 75 × 25 mm substrate slides (glass, tile [i.e. glazed ceramic], plastic [i.e. acryl], and wood [i.e. untreated pine]) were inserted into PVC pipe sections using polyethylene foam and ethyl cyanoacrylate. The substrates and their holders were then either sewn into 25.4 × 30.5 cm 100 µm mesh enclosures to protect from predation vie exclusion or left exposed to the native environment (Supplementary Figure 1).

Substrates were submerged in the Otago Harbour, New Zealand (45.826678 S, 170.641684 E). Samples were suspended from a rope using cable ties 80 cm from the seabed, remaining exposed during low-tide but submerged during high-tide. All samples were located in a small geographic area (< 30 m^2^), 50 metres from shore.

Starting in May 2019 triplicate biofilm sampling was completed on days 7, 14, 19, 28, 42, and 56. However, due to sample loss as a result of the *in situ* environment, a total of 153 biofilm samples were obtained (Supplementary Table 1). Biofilm was scraped off the entire substrate using a sterile scalpel. Tiles were scraped only on the glazed side. Samples were suspended in 100 µL sterile Milli-Q water and stored at - 80 °C until further processing.

**Table 1.** ANOSIM and ADONIS tests compare time, substrate and enclosure correlations over time on biofilm prokaryotes and eukaryotes.

| Organism | Test | Biofilm development stage | Variable | R/R<sup>2</sup> | p value |
| --- | --- | --- | --- | --- | --- |
| Prokaryotes | Anosim | Early | Time | 0.3 | < 0.01 |
|  |  |  | Substrate | 0.5 | < 0.01 |
|  |  |  | Enclosure status | 0.19 | < 0.01 |
|  |  | Late | Time | 0.3 | < 0.01 |
|  |  |  | Substrate | 0.19 | < 0.01 |
|  |  |  | Enclosure status | 0.41 | < 0.01 |
|  | Adonis | Early | Time | 0.11 | < 0.01 |
|  |  |  | Substrate | 0.24 | < 0.01 |
|  |  |  | Enclosure status | 0.08 | < 0.01 |
|  |  | Late | Time | 0.13 | < 0.01 |
|  |  |  | Substrate | 0.12 | < 0.01 |
|  |  |  | Enclosure status | 0.13 | < 0.01 |
| Eukaryotes | Anosim | Early | Time | NA |  |
|  |  |  | Substrate | 0.34 | < 0.01 |
|  |  |  | Enclosure status | 0.52 | < 0.01 |
|  |  | Late | Time | 0.32 | < 0.01 |
|  |  |  | Substrate | 0.06 | < 0.01 |
|  |  |  | Enclosure status | 0.23 | < 0.01 |
|  | Adonis | Early | Time | NA |  |
|  |  |  | Substrate | 0.28 | 0.014 |
|  |  |  | Enclosure status | 0.18 | < 0.01 |
|  |  | Late | Time | 0.16 | < 0.01 |
|  |  |  | Substrate | 0.05 | 0.018 |
|  |  |  | Enclosure status | 0.07 | < 0.01 |

Duplicate 1L water samples were collected 2 metres from both ends of the substrate suspension structure on days 0, 7, 14, 19, 28, 42, and 56. Samples were filtered through a 0.22 μm (diameter = 47 mm) polycarbonate filter prior to freezing and storage at -80 °C until further processing.

All data analysis was carried out using R version 3.6.1 within RStudio ^71^, and visualised using the ggplot2 package (version 3.2.1) ^72^ unless otherwise stated. All code and associated files are available at https://github.com/SvenTobias-Hunefeldt/Biofilm_2020/.

### DNA Extraction and Sequencing

DNA was extracted using the Qiagen DNeasy® PowerSoil® Kit (MoBio Laboratories, Carlsbad, CA, USA) according to the manufacturer’s protocol. DNA was then dehydrated using rotary evaporation in an Eppendorf Concentrator 5301(Eppendorf, Germany) at 30°C for 1 hour. Community profiles were generated using barcoded 16S (targeting the V4 region: 515F [5′-NNNNNNNNGTGTGCCAGCMGCCGCGGTAA-3′] and 806R [5′-GGACTACHVGGGTWTCTAAT-3′]) or 18S (1391f [5′-GTACACACCGCCCGTC-3′] and EukBr [5′-TGATCCTTCTGCAGGTTCACC TAC-3′]) rRNA gene primers as per the Earth Microbiome Project protocol ^73^. Barcoded samples were then loaded onto separate Illumina MiSeq 2 × 151 bp runs (Illumina, Inc., CA, USA) to produce a total of 13,202,642 and 10,124,722 reads for 16S and 18S runs, with an average of 63,171 and 48,444 per sample and a standard deviation of 28,248 and 18,473. All sequence data from this study has been deposited in NCBI under BioProject PRJNA630803.

16S and 18S rRNA sequencing reads were quality filtered and assigned to amplicon sequencing variants (ASVs) using the dada2 R package (version 1.12.1) and associated pipeline ^74^. Taxonomies were assigned in accordance to the dada2 pipeline from the SILVA rRNA reference database (version 132) using the Ribosomal Database Project (RDP) naïve Bayesian classifier method ^75^. All data was imported into R for further analysis with the phyloseq R package (version 1.28.0) ^76^.

The optimum sequencing depth was identified with a custom function utilising functions from the phyloseq, reshape2 (version 1.4.3) ^77^, plyr (version 1.8.4) ^78^, base (version 3.6.1) ^71^, stats (version 3.6.1) ^71^, and data.table packages (version 1.12.1) ^79^. Samples were rarefied 10 times at 11 000 reads using rarefy_even_depth() from the phyloseq package. This depth retains both the maximum number of samples and sequencing depth (Supplementary Figure 2). Independent rarefactions were combined and underwent sample count transformations using transform_sample_counts() from the phyloseq package to account for multiple rarefactions. To avoid fractional representation of counts all data was rounded to the nearest possibility using the round() command from the stats package. All data analysis used rarefied data unless otherwise stated.

### Community analysis

Observed richness was quantified using estimate_richness() from the phyloseq package, and significance analysed using stat_compare_means() from the ggpubr package (version 0.2.4) ^80^. Phyloseq package generated NMDS plots assessed Beta-diversity, with ggplot2 and ggpubr package adjustments. Clustering was assessed using ADONIS and ANOSIM statistical tests, with the vegan package (version 2.5-6) ^81^, pairwise PERMANOVA tests identifying significant sample to sample differences. Spearman correlations were used to assess relationships between biofilm age and NMDS1 of the whole community NMDS. Intra-time dissimilarity was calculated with the vegan and phyloseq package, removing self-comparison using the dplyr package (version 0.8.3) ^82^. Significant differences between time-points were identified with the stats package. The number of shared organisms was calculated using Zeta.decline.mc() from the zetadiv package (version 1.1.1) ^83^.

Comparisons of means between two groups, such as biofilm succession stages and predated conditions, were done with Wilcoxon tests. In the case of more than two groups Kruskal-Wallis tests were used, e.g. differences between biofilm ages and substrates. Pairwise Wilcoxon tests compared individual timepoints and substrates.

To test for taxonomic composition changes we determined what organisms significantly changed over time using the EdgeR package (version 3.22.3) ^84^ and an exact test using only biofilm samples. The legend was ordered according to mean relative abundance with the forcats package (version 0.4.0) ^85^. All p values were FDR adjusted using Bonferroni. A literature search was used to classify phyla as either autotrophs ^86–89^, heterotrophs ^51,60,61,90–98^, mixotrophs ^90,91,99,100^, or unknown if the literature was lacking. Unknown taxonomies were excluded due to their low abundance (Supplementary Table 1), and unknown role in predator-prey dynamics.

Rare taxa (phyla making up less than 1% of the total abundance) were identified using the plyr package and standard error, standard deviation and the 95% confidence interval calculated with the Rmisc and dplyr packages.

## Results

### Biofilm maturation is divided into two stages

Biofilms across all treatments could be separated into two distinct succession stages: early and late. Silhouette analyses identified that prokaryotes were best divided into 2 groups, predominantly based on community age, whereas eukaryotic groupings could not be reliably identified. Ecotone analyses found only 2 (day 7 and 14 for prokaryotes) and 1 (day 7 for eukaryotes) out of 6 time-points categorized as early stage (Supplementary Figure 3). Richness and compositional patterns were in agreement with the rapid successions. Richness increased with biofilm age (Kruskal-Wallis, chi^2^ > 19, p < 0.01) with a late-stage 1.9-fold increase compared to the early-stage (Figure 1 A-B) (Wilcoxon, chi > 407, p < 0.01). Pairwise analyses further confirmed differences between succession stages (Wilcoxon p < 0.03), with non-significant intra-stage differences (Wilcoxon p > 0.06). Compositional patterns also exhibited stage significant grouping, both when comparing group means (ANOSIM, R = 0.67, p < 0.01) and when taking variability into account (ADONIS, R^2^ = 0.13, p < 0.01). Additionally, community dissimilarity displayed stabilisation from day 19 (prokaryotes) and 14 (eukaryotes) to 42 (Figure 1 C), with early-stage samples sharing less ASVs (Amplicon Sequence Variants) across substrates (Figure 1 D).

**Figure 1.**
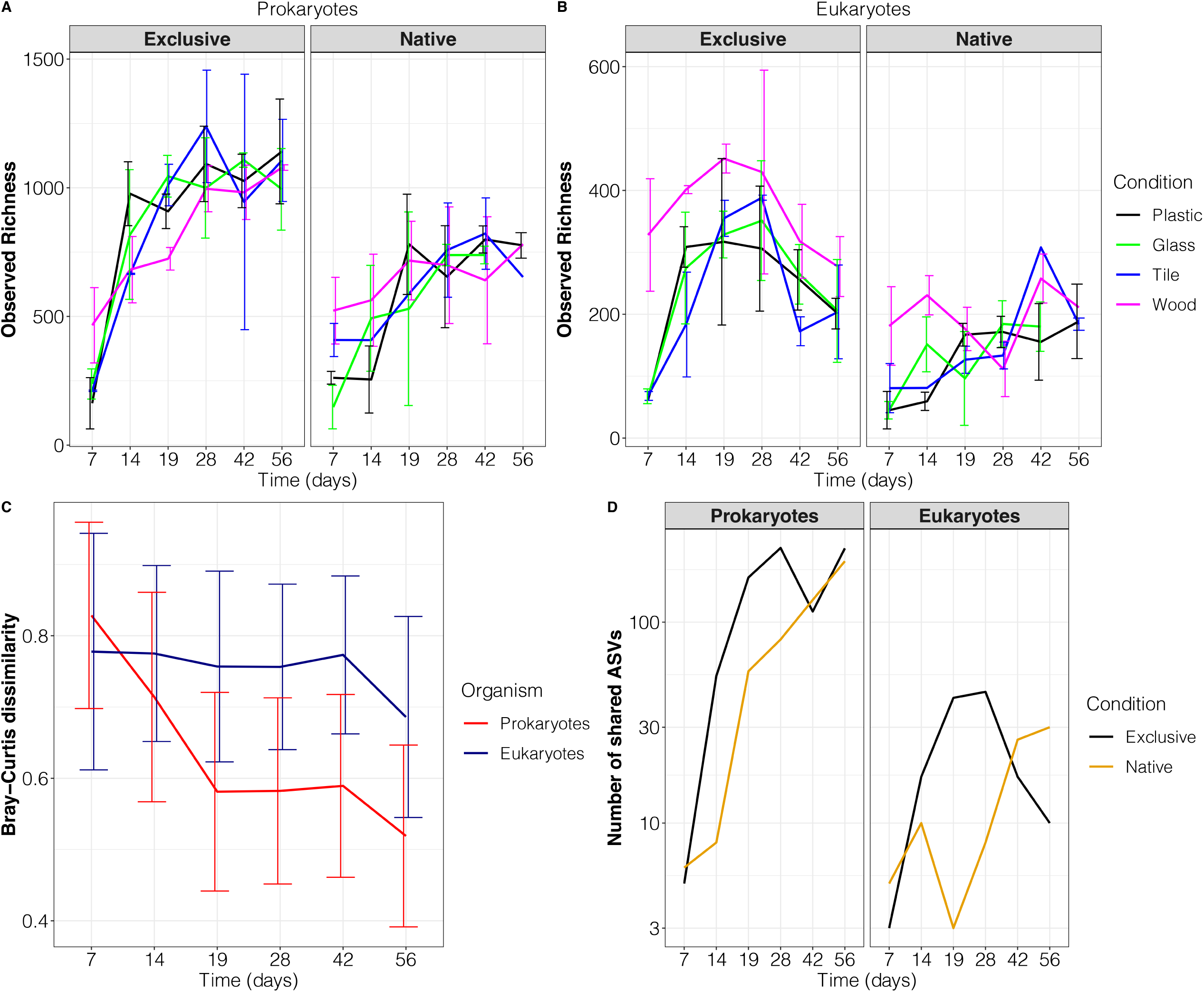
Marine biofilm development occurs in two stages. Community richness changes over time are seen for both prokaryotic (A) and eukaryotic (B) organisms. Also shown is the intra-time point community dissimilarity of each biofilm (C) and the shared number of ASVs (Amplicons Sequence Variants) between substrates over time (D). Colours depict both different substrates (A & B), organisms (C), and enclosure conditions (D). Black, green, blue, and magenta represent plastic, glass, tile, and wood (A & B), while red and blue represent prokaryotic and eukaryotic organisms (C), and black and orange represent exclusive (enclosed) and native (non-enclosed) conditions (D). Error bars represent standard deviation.

#### Rapid successions lead to community convergence across substrates

Rapid community successions were observed with sequential clustering in response to biofilm age (Table 1, Figure 2). Biofilm age also significantly correlated with NMDS1 (Figure 2) (Spearman, rho > 0.66, p < 0.01), and both maturation and sequential clustering coincided with community convergence across substrates. Variance between substrates and enclosure status decreased over time, especially when comparing specific time points between early and late community stages, such as day 7 and 56 (Figure 1 C). However, prokaryotic variance decreases were more pronounced than for eukaryotes (28% and 9% across the experiment for prokaryotes and eukaryotes, respectively) (Figure 1 C). Both prokaryotes and eukaryotes showed increased ASV sharing across substrates over time (Figure 1 D). The number of shared ASVs across substrates increased from 1.8 (prokaryotes) and 3.3% (eukaryotes) to 22 and 9.4% respectively of the total number of community ASVs based on richness. Predator exclusion increased the rate of prokaryotic community maturation (this difference could be quantified as a dissimilarity increase of 0.02% per day). Over 56 days the difference in community succession velocities lead to a maximum dissimilarity of 11.2%, however, day 56 prokaryotic communities are a mean 61% dissimilar. Community convergence was associated with a transition from bottom up to top down control. Early clustering was strongest due to substrate choice (Table 1, Figure 2), with a distinct wood composition (pairwise MANOVA, p < 0.03), and no observable early stage distinctions between enclosure status (Supplementary Figure 4). Meanwhile late communities clustered primarily due to enclosure differences (Table 1, Supplementary Figure 4) with minimal substrate dependent grouping (Figure 2).

**Figure 2.**
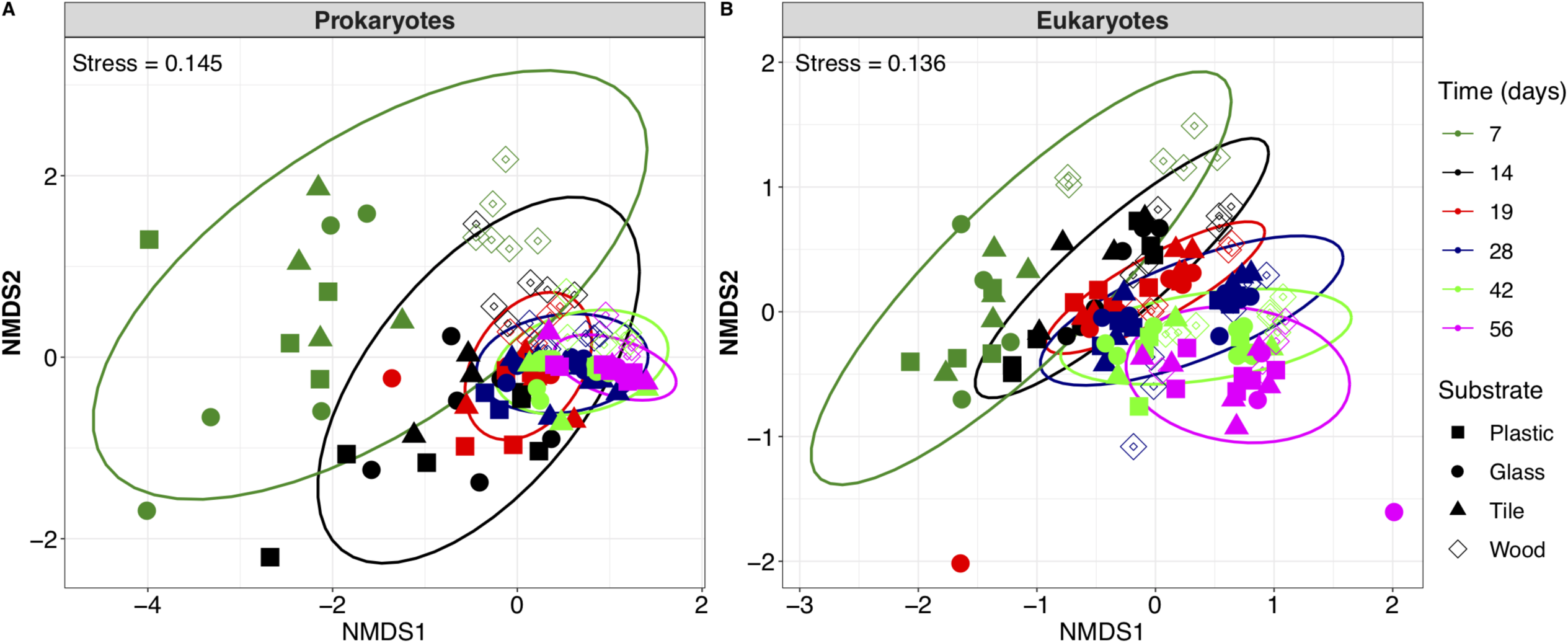
Microbial biofilm beta-diversity by time. NMDS plots show prokaryotes (A) and eukaryotes (B), ellipses depict the 95% confidence interval grouping effects of time with biofilm age represented as different colours (days 7, 14, 19, 28, 42, and 56 depicted as olive, black, red, navy, green, and magenta). Substrates are depicted with different shapes, the hollow diamond represents wood whereas the filled square, circle, and triangle represent plastic, glass, and tile. Stress is included in the upper left corner of individual plots.

#### Native conditions displayed decreased alpha diversity, with some substrate effects

Predator exclusion exerted noticeably more pressure on late than early communities. Exclusion increased late stage richness by 2-fold (Wilcoxon p value < 0.01) (Figure 3, Figure 1 A-B) while early communities remained unaffected (Wilcoxon p > 0.06). Substrate-specific differences were enclosure-dependent: 100-250 more organisms were detected within early native prokaryotic and late exclusive eukaryotic wood communities compared to more inert substrates (Figure 3). On average 42 less prokaryotic ASVs were detected on wood compared to inert substrates, while eukaryotes had a mean of 79 more ASVs on wood (Figure 3). However, prokaryotic substrate richness differences were only significant when wood contained an increased number of ASVs compared to inert substrates, such as under native early conditions. Overall, richness was primarily driven by enclosure condition, with some substrate dependent differences.

**Figure 3.**
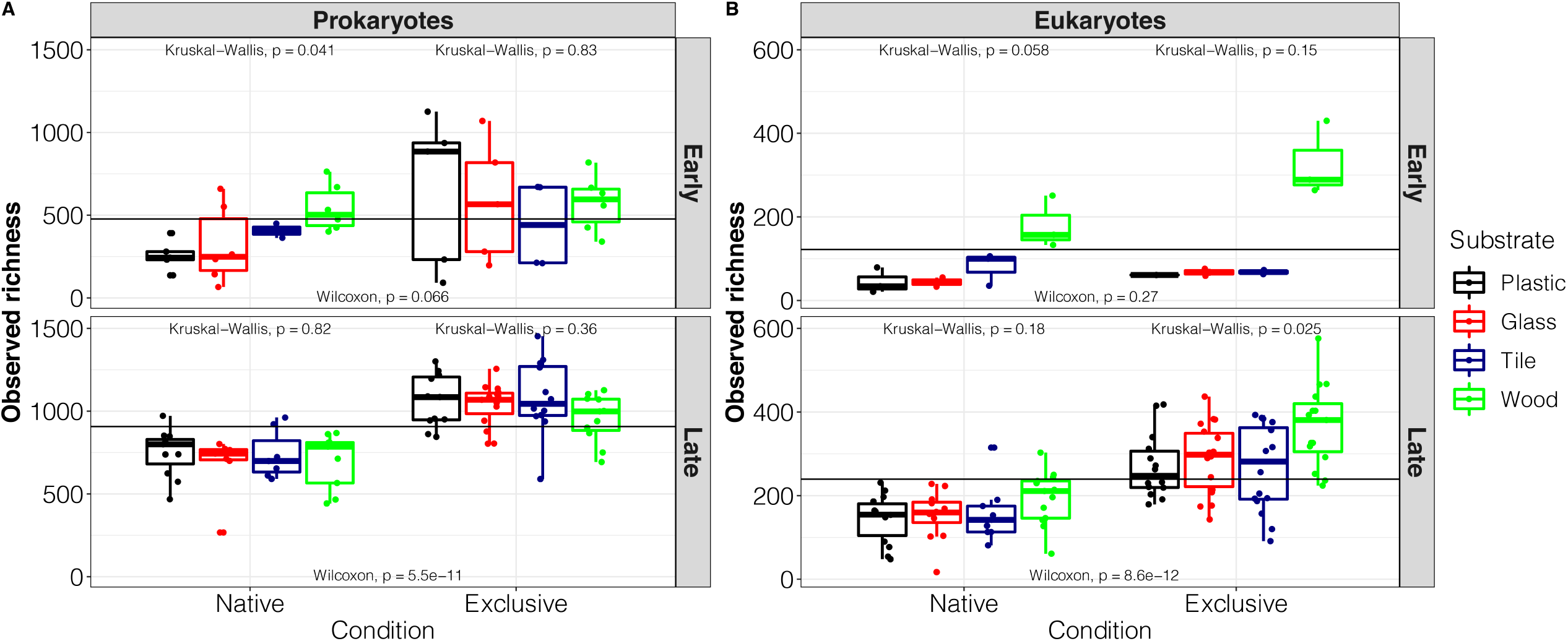
Prokaryotic and eukaryotic observed richness is enclosure specific in a stage dependent manner. Marine *in situ* biofilm species richness for both prokaryotes (A) and eukaryotes (B) is shown by developmental stage grouping. The black horizontal line represents the organisms and stage specific richness mean, while colours separate the different substrates. Plastic, glass, tile, and wood is depicted in black, red, navy, and green. Wilcoxon tests assessed the significance of stage specific enclosure differences, and Kruskal-Wallis tests identified the significance of substrate effects in a stage and enclosure specific manner.

### Shifts in dominant taxa reflect changes in selective pressures

Phylum level changes were observed in response to switch in bottom-up to top-down control mechanisms (Figure 4). Early communities differed primarily by substrates; wood displayed 9-50% more Proteobacteria and Euglenozoa and half the number of autotrophs compared to inert substrates (Supplementary Table 2). The absence of an enclosure caused the main difference in late composition which displayed no consistent substrate effect. Enclosed biofilms were dominated by mixotrophs and heterotrophs while native conditions contained high autotroph values. Late enclosed community autotroph relative abundance was half of the non-enclosed conditions. Ochrophyta abundance, predominantly diatoms, decreased from > 36% to ∼15% across all substrates. Meanwhile, Cyanobacteria abundance increased in glass, tile, and acyl plastic from <30% to >53% from days 7 to 14, and then decreased to ∼17%, whereas wood Cyanobacteria abundance rose by 10%. However, with autotroph decreases came mixotroph and heterotroph increases. Over 56 days Proteobacteria abundance increased by a mean 16.7% while Bacteroidetes, Ciliophora, and Arthropoda abundance rose by 11%, 21%, and 49% (Supplementary Table 2). Within non-enclosed biofilms Bacteroidetes, Ciliophora, and Arthropoda stayed < 25% abundance, while Cyanobacteria increased by 21% within wood and remained > 45% within all other substrates. Early autotroph-heterotroph balances are therefore due to bottom-up substrate differences. Followed by a primary selective pressure switch, that decreased early community differences, through decreases of heterotroph previously abundant on wood until late autotroph-heterotroph distributions were enclosure dependent.

**Figure 4.**
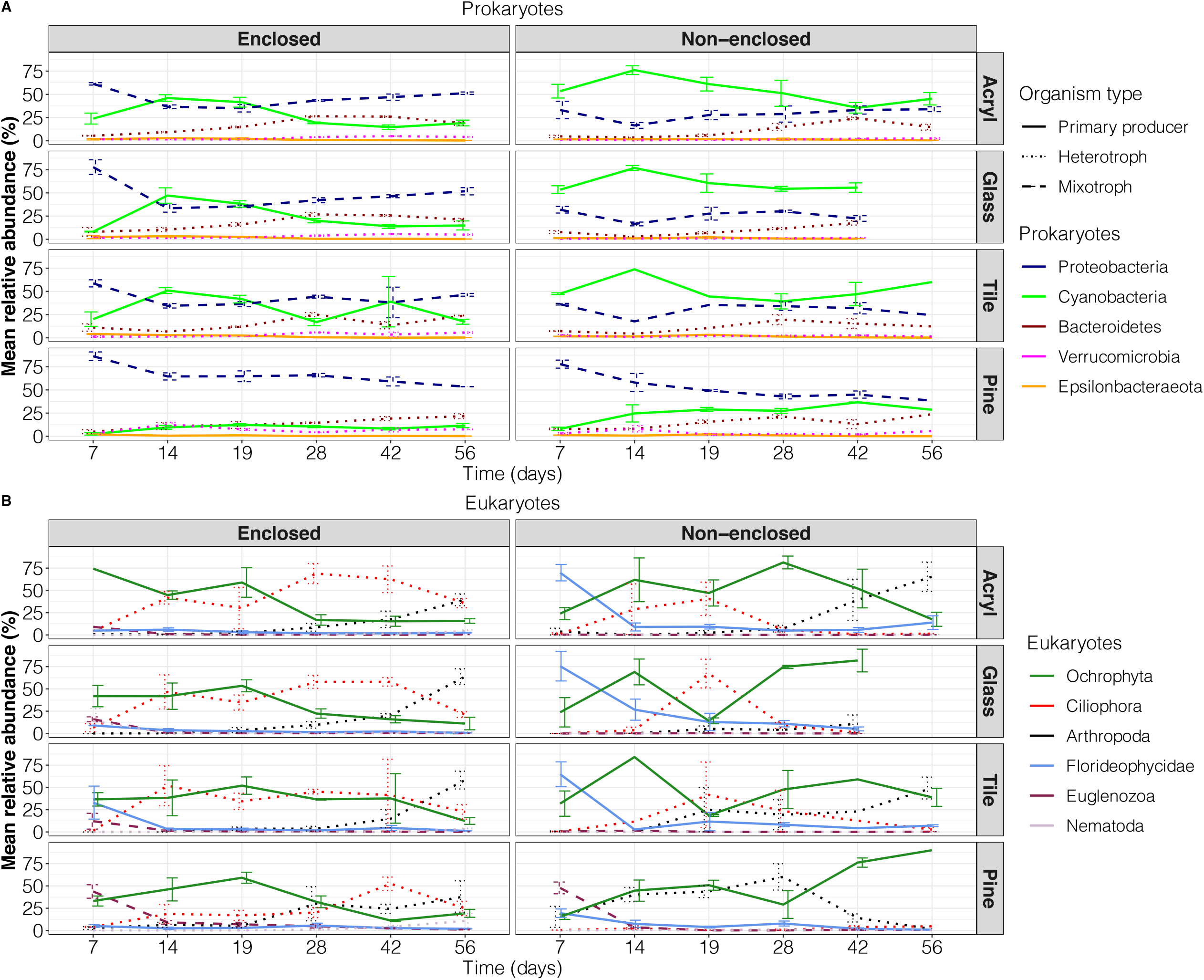
Significant phylum changes in response to biofilm age. The mean relative abundance of significantly prokaryotic (A) and eukaryotic (B) phyla correlated with biofilm age are depicted with exclusive (enclosed) communities on the left and native (non-enclosed) on the right in a substrate dependent manner. The type of line (solid, points, and dashed) represent the three main types of organisms; autotrophs, heterotrophs, and mixotrophs. Mean relative abundance was calculated from pooled significantly correlated taxa and error bars represent the standard error of the mean abundance based on biological replicates.

## Discussion

Our results show that the analysis of bottom-up selective pressures (i.e. substrate differences) are sufficient to determine the majority of community variance when assessing drivers of early unstable communities, but the study of top-down selective pressures (i.e. predation via a mesh enclosure) is essential when assessing assembly mechanisms of established environments as it determines the majority of community variance. This supports the hypothesis that early communities are primarily driven by bottom-up controls but did not support our prediction for late communities. Late taxonomic and diversity patterns were instead predominantly determined by top-down control. From our results, we created a community assembly model (Figure 5) describing the effect of sequential bottom-up and top-down control over time. Initial community settlement is a stochastic process that results in large variation. Subsequent growth is then determined primarily by nutritional availability and results in distinct compositions based on dominant factors like availability and type of nutrients (i.e. degradable vs non-degradable substrates). Early bottom-up control weakens over time until late compositions are modified by top-down selective pressures. This work addresses some of the controversy in the field of ecology, by quantifying the relative effects of bottom-up and top-down selective pressures over time. This approach also assesses the effects of top-down pressures on community composition and assembly in more complex environments, which represents a gap in the literature ^37,38^. Below we discuss the transition from a stochastically settled to a top-down controlled community via convergent selection, as well as the effects of substrate properties and predation on compositional patterns.

**Figure 5.**
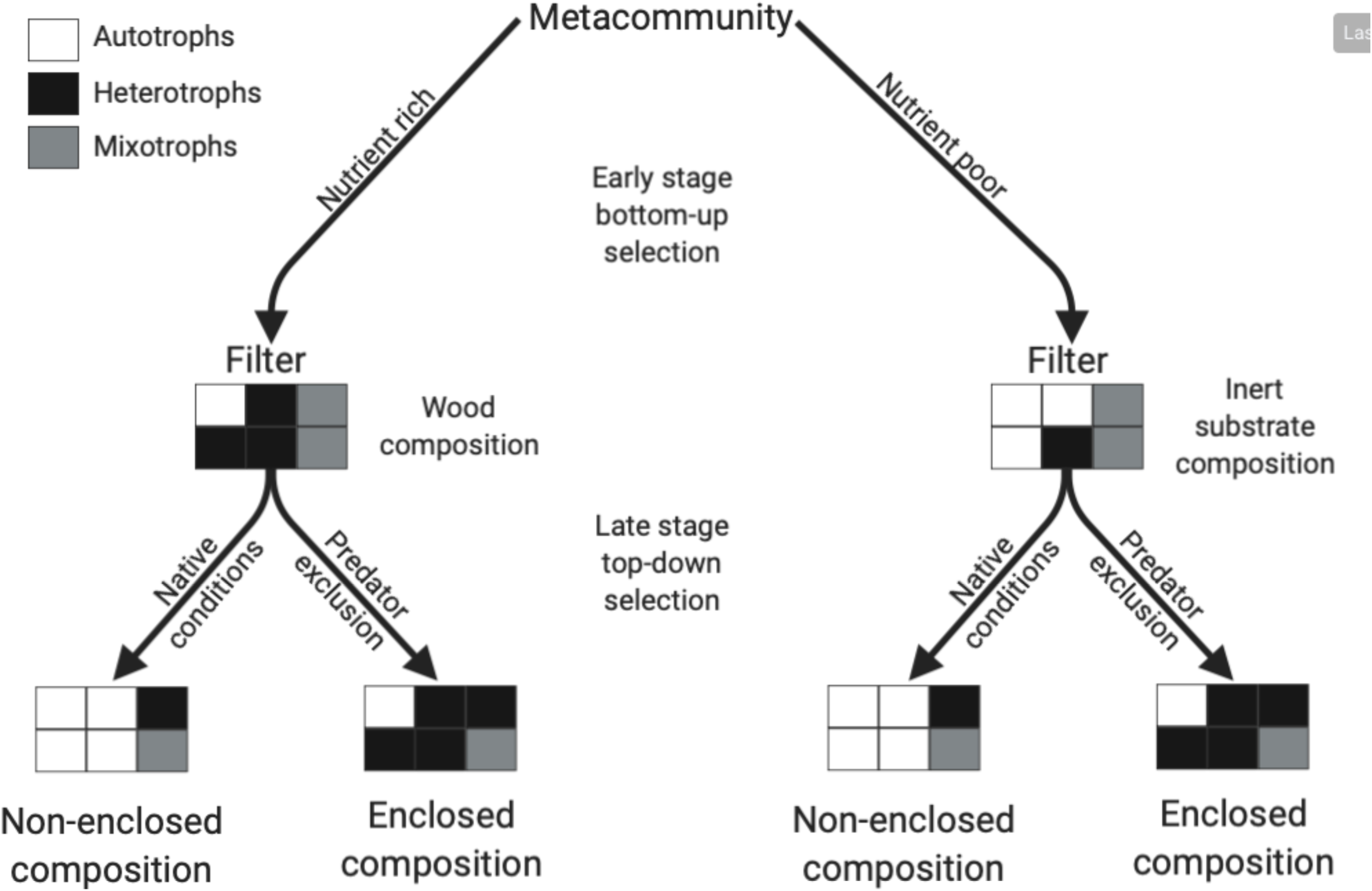
Microbial biofilm community assembly is divided into discrete stages associated with distinct compositions and selective pressures. Communities are a subset from the environmental organism pool known as the metacommunity and filtered in response to a selective pressure. Selective pressures filter for distinct community compositions. First, bottom-up (early stage) substrate differences and then top-down (late stage) enclosure status determine biofilm community composition, with decreased bottom-up differences due to top-down filtering.

### Stochastic settlement transitions to convergent community assembly

Settlement is primarily a stochastic process, while subsequent deterministic processes result in convergence. Previous studies have found that dispersal and selection influence composition ^7,8^. The limited geographical range of our study controls for unequal dispersal, so encounter rates dictated early composition. Large early community confidence intervals and intra-age dissimilarities thereby reflect the inherent stochasticity of early-stage selection. Previous studies have found that early assembly is dictated predominantly by stochastic mechanisms ^39^. However, deterministic processes may still play a role in settlement. Early settlers must adhere to the substrate surface for colonisation, and a mesh enclosure excluded organisms over 100 µm. Yet no enclosure-specific visual clustering was identified within early biofilms. A size exclusion method was chosen as predators tend to be larger than their prey ^40^. Previous work has also shown that larger organisms (e.g. macro-algae and macro-eukaryotes) are not primary settlers, and instead prefer established and complex communities ^41^. Therefore, while surface attachment was a critical process, the enclosure size exclusion had no effect on settlement.

Following settlement, communities can develop either divergently ^42–44^ or convergently ^5,43,45^, with evidence for both rife within the literature. Stochastic fluctuations throughout time and the creation of new genetic variation, drift and diversification, are characteristic of time-dependent ecological studies ^7^ and lead to divergence with increased dissimilarity over time. Our study however displayed convergence via an increased number of shared organisms over time and decreased dissimilarity. The rise of convergence is most likely an artefact of preconditioning ^46^ due to the geographically-limited environment and therefore decreased organism variability, and the surface adhesion requirement. Another point in favour of convergence is derived from deterministic succession ^12^. Community age correlated with the primary axis of dissimilarity (NMDS1) and clustered in response to biofilm age. These characteristics imply succession due to sequential compositional changes, with the same organisms filling identical ecological niches. Based on the evidence we conclude that compositions stabilised over time, with unstable early communities while late communities were more stable due to community convergence. The transition from unstable to stable compositions is seen in both developing and disrupted communities as they undergo rapid changes as ecological niches develop and fill in response to rapid condition changes. These rapid changes therefore cause disturbed communities to ‘reset’ ^47^. As such, our observations regarding early communities (day 7-14 within prokaryotes and day 7 within eukaryotes) are applicable to both new and recently disturbed communities.

### Substrate differences drive early composition

Increased degradability and subsequent higher nutrient availability and roughness of wood could be responsible for higher richness in early predated conditions. Grooves within rough surfaces increase the degree of colonisation through increased surface area, and protection from shear stress and large predators ^48^. Increased richness is therefore expected of rougher surfaces such as wood ^49,50^. However, wood also displayed a significantly dissimilar community compared to other substrates under both exclusive and native conditions. The organic carbon within wood in the form of lignocellulose is accessible to mixotrophic and heterotrophic lignocellulose degraders (i.e. *Proteobacteria* and *Bacteroidetes* ^51,52^), thereby the organic carbon could act as an environmental filter. Inert substrates (i.e. plastic, glass, and tile) cannot be degraded in the same manner, as such their early community was instead primarily composed of autotrophs which fix their own carbon. Wood’s organic carbon availability therefore shapes early composition through the promotion of heterotrophic growth.

We hypothesise that substrate dependent differences decrease over time due to increased community density. Previous studies have shown that increased density leads to biofilm heterogeneity ^53^ as organisms further removed from the nutritional supply are affected by solute diffusion limits ^53^. In the context of our study, organisms further away from the substrate surface cannot utilise wood’s nutritional organic carbon. Increased substrate coverage and subsequent displacement from the substrate surface decreased differences between degradable and inert substrates over time. We conclude that substrate dependent differences decrease over time due to increased density limiting surface derived nutritional supplies and protection to cause convergence. As seen in the similar community richness and composition in late biofilms regardless of substrate type.

### Predation controls diversity and established ecosystem composition

Predation decreased richness, although differences between substrates remained. The effects of predation on microbial richness are relatively unexplored. Predation has been shown to increase or decrease community richness, ^54,55^ dependent on the net number of directly or indirectly repressed/enriched organisms ^56^. Our study determined that predation decreases diversity. While substrate diversity effects could be identified, they remained mainly predation-specific differences between substrates, e.g. wood showed increased richness within early predated prokaryotes and late enclosed eukaryotes.

Predation affected late established communities to a greater extent than their early counterparts. Late communities exhibited qualities associated with established ecosystems ^57,58^, via decreased richness and dissimilarity variability, indicative of increased community stability. Early communities lack the inherent complexity and resources to sustain large abundances of predators and other heterotrophic organisms, unlike their late counterparts ^41^. Predation significantly decreased late richness, but early richness remained unaffected. Heterotrophs such as Arthropoda and Ciliophora increased in abundance over time and drove their prey below detectable limits. Arthropoda prey on all microbial life ^59,60^, whereas Bacteroidetes prefer particulate detritus such as dead autotrophic Ochrophyta, while also being capable of surface degradation ^51,61^. Heterotrophic organism increases mirrored autotroph decreases, specifically Cyanobacteria and Ochrophyta. Predated conditions display the autotroph to heterotroph shift over time as higher trophic levels preferentially preyed upon heterotrophs.

Heterotrophs have been predicted to be more nutritious and thus a more important energy source compared to autotrophs. Autotrophs display weak homeostasis, with stoichiometric ratios shaped by the environment to a larger degree than the strongly homeostatic heterotrophs ^62^. Additionally, the environment, and therefore autotrophs, contains a higher carbon to phosphorous ratio compared to heterotrophs ^62,63^. The predation of stable and closer stoichiometric matches leads to easier heterotrophic stoichiometric ratio maintenance, making heterotrophs a better nutrient source compared to autotrophs. The closer stoichiometric matches could explain the closer relationships between heterotrophs and higher trophic levels ^64^, such as the inverse relationship between Ciliophora (a heterotroph) and Arthropoda (a eukaryotic predator). Previous findings corroborate the relationship: Ciliophora (Ciliates) abundance declines corresponding with increases in Arthropoda (Copepods), with no alternate prey availability effects ^65^. Overall, we found that these taxonomic responses confirm that predation is an important driver of late composition.

Previous work was completed within controlled environments with limited numbers of external inputs, as opposed to the more complex and dynamic habitat approach of our study^37^. However, both previous and this study’s findings agree that higher-order biotic interactions play a critical role in complex microbiomes^37^. Often leading to community convergence and stabilisation.

Alternative interpretation to our results are possible due to physical impacts of enclosure. Enclosed conditions decrease the flow rate dependent nutrient supply and shear stress, as well as light availability. A decreased organic nutrient supply leads to higher autotroph abundances as other organisms lack essential nutrients to fuel their growth and reproduction. Enclosed conditions should therefore display higher autotroph abundances, but this was not the case. Our study instead showed that autotrophs (i.e. Cyanobacteria, Florideophycidae, and Ochrophyta) were, as a general rule, decreased within enclosed conditions. This could be due to light availability limitations when over time more organisms settle on the mesh enclosure and decrease light penetration, as well as the mesh itself providing shade ^66,67^. Based on this a linear relationship between autotrophs and time under enclosed conditions would be expected, as more light is blocked with increased mesh community density. Instead Cyanobacteria and Ochrophyta relative abundances under enclosed conditions peak at day 14 and 19, suggesting that light availability due to growth on the mesh may not be a large contributing factor compared to other compositional drivers, while the inherent shade effect of mesh may still play a role. Strong mesh inherent shade effects would lead to large enclosure status effects throughout the study, due to large autotroph decreases within exclusive communities. However, early community differences were primarily dependent on substrate surface properties. Late community differences may be due to the mesh shade effects, but this is also unlikely as enclosure status independent predation dynamics increased over time, such as the close inverse relationship between Ochrophyta and Ciliophora. Therefore, we conclude that community compositions were predominantly due to substrate and predation effects, rather than the consistent effects of an inherent mesh shade. We also considered that the presence of an enclosure may lead to community differences through sheer stress mitigation ^68,69^, but thought it unlikely due to the following reasons. First, shear stress acts through biofilm biomass removal ^70^ and should thereby equally affect all organisms rather than selectively removing one group, so relative abundances would remain the same. Second, a lack of shear stress would increase settlement, but no significant richness difference exists between early enclosure conditions. Third, high stress has been shown to decrease biofilm maturation rates ^68,69^, but the majority of compositional differences between enclosures in our study was not due to maturation rate variations. Therefore enclosure differences are more likely due to predator effects rather than shear stress removing biomass and decreasing maturation rates.

### Conclusion

Within our study we identified a switch from bottom-up to top-down control linked to community maturation. This switch in primary selective pressure highlights the community selection of primary colonisers, to the emergence of stable and convergent mature communities modified by predation (Figure 5). Predation selectively removes organisms, potentially using nutritional value as a criteria. The integration of top-down controls allowed for the explanation of more variance than a pure solely bottom-up focus. Hence, increased importance should be placed on top-down controls, particularly for studies on stable late communities. The integration of both bottom-up and top-down selective pressures in the field of microbial ecology leads to increased knowledge of assembly mechanism. This knowledge would be applicable across all ecosystems, as the underlying framework rather than habitat specific processes, is assessed. Ultimately identifying not only habitat specific community compositions, but what lead to these compositional patterns.

## Acknowledgements

We thank Dave Wilson for his contribution to experimental set up.

## Data availability

The sequence data from this study have been deposited in NCBI under BioProject PRJNA630803. All data generated and/or analysed during the study is available within the GitHub repository, https://github.com/SvenTobias-Hunefeldt/Biofilm_2020/.

## Figure Legends

**Supplementary Figure 1. Biofilm holder construction and deployment**. Individual 75 × 25 mm substrate (i.e. plastic, glass, tile, and wood) slides were contained within 50 mm PVC pipe sections with a diameter of 100 mm with the use of polyethylene foam and ethyl cyanoacrylate. Predators were excluded with the use of a 100 µm mesh surrounding the substrate filled PVC pipe section, whereas native (non-enclosed) samples lacked a mesh enclosure. Samples were suspended from either their mesh enclosure or pipe on a single 30 m rope on the surface during low tide 80 cm from the seabed, so high tide submerged samples.

**Supplementary Figure 2. Prokaryotic and eukaryotic organism rarefaction curves**. Mean observed richness was identified in response to sampling depth for prokaryotes (A) and eukaryotes (B), with 10 replicate rarefaction and richness determination events. Different colours represent biofilm age, and error bars represent the standard deviation of individual sample replicates.

**Supplementary Figure 3. Development stage evidence based on a silhouette and ecotone analysis for prokaryotes (left) and eukaryotes (right)**. Determination of the ecotone transition zone (A, B) and a silhouette analysis (C, D) was carried out for both prokaryotic (A, C) and eukaryotic (B, D) organisms. The ecotone analysis was done using biofilm age as a distance proxy to facilitate analysis. Colours represent the two identified groups with the average species abundance and biofilm age depicted on the y axis and x axis.

**Supplementary Figure 4. Development stage depicted beta-diversity**. NMDS plots show prokaryotic (A) and eukaryotic (B) organism beta-diversity, colours separate the early (black) and late (red) succession stages, surrounded by 95% confidence interval ellipses. The enclosure status is depicted with different shapes, the hollow diamond represents native (non-enclosed) samples whereas the filled circle represents exclusive (enclosed) samples.

**Supplementary Table 1**. Sampling replicates from Dunedin Harbour over the course of the study, based on time of sampling, and community source.

**Supplementary Table 2. Significant taxonomic changes over time**. Significant relative taxonomic (i.e. family) changes in response to biofilm age separated by substrate and enclosure status.

