## Supplementary figures and images for "Predation impacts late but not early community assembly in model marine biofilms"

### Supplemental Figure S1

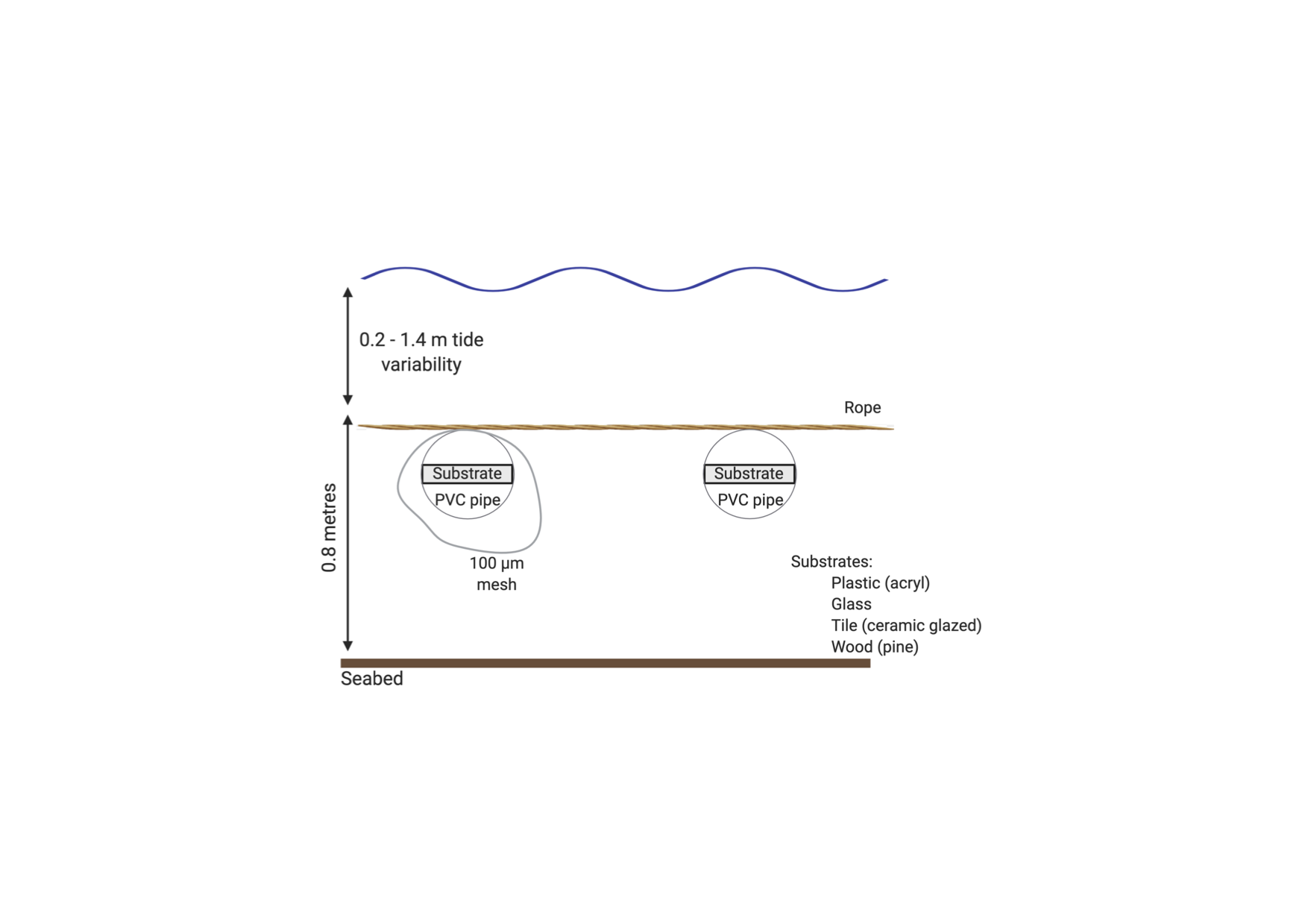

### Supplemental Figure S2

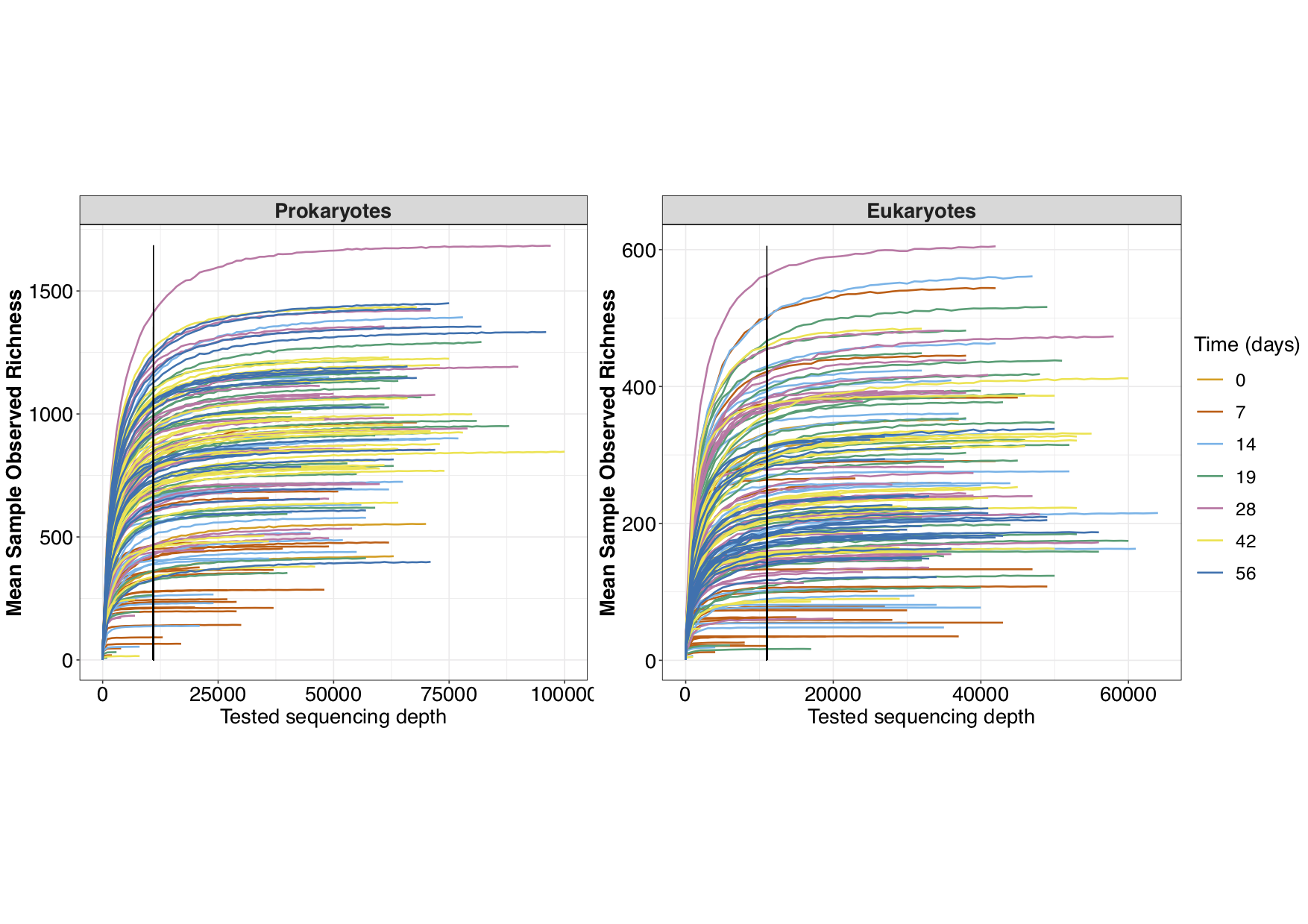

### Supplemental Figure S3

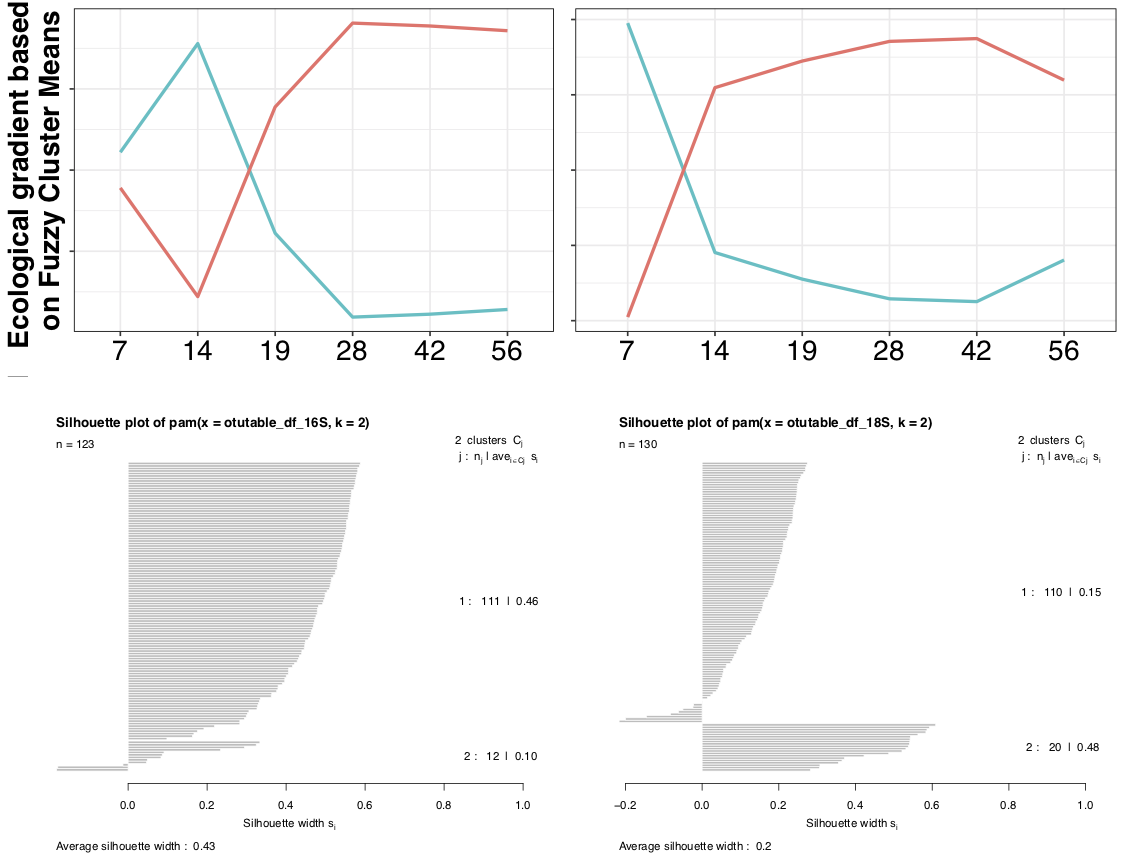

### Supplemental Figure S4

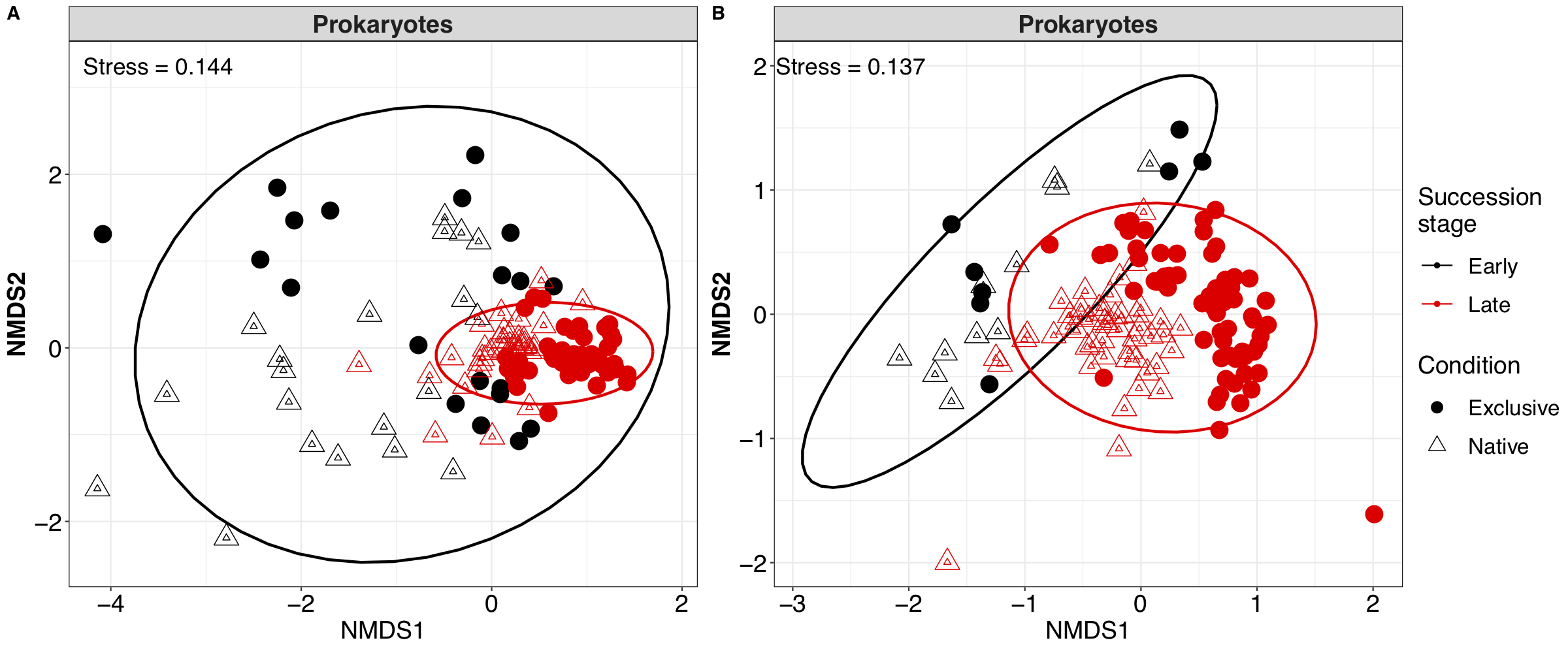
